# Ancient genomic variation underlies repeated ecological adaptation in young stickleback populations

**DOI:** 10.1101/167981

**Authors:** Thomas C. Nelson, William A. Cresko

## Abstract

Adaptation in the wild often involves standing genetic variation (SGV), which allows rapid responses to selection on ecological timescales. However, we still know little about how the evolutionary histories and genomic distributions of SGV influence local adaptation in natural populations. Here, we address this knowledge gap using the threespine stickleback fish (*Gasterosteus aculeatus*) as a model. We extend the popular restriction site-associated DNA sequencing (RAD-seq) method to produce phased haplotypes approaching 700 base pairs (bp) in length at each of over 50,000 loci across the stickleback genome. Parallel adaptation in two geographically isolated freshwater pond populations consistently involved fixation of haplotypes that are identical-by-descent. In these same genomic regions, sequence divergence between marine and freshwater stickleback, as measured by *d*_*XY*_, reaches ten-fold higher than background levels and structures genomic variation into distinct marine and freshwater haplogroups. By combining this dataset with a *de novo* genome assembly of a related species, the ninespine stickleback (*Pungitius pungitius*), we find that this habitat-associated divergent variation averages six million years old, nearly twice the genome-wide average. The genomic variation that is involved in recent and rapid local adaptation in stickleback has actually been evolving throughout the 15-million-year history since the two species lineages split. This long history of genomic divergence has maintained large genomic regions of ancient ancestry that include multiple chromosomal inversions and extensive linked variation. These discoveries of ancient genetic variation spread broadly across the genome in stickleback demonstrate how selection on ecological timescales is a result of genome evolution over geological timescales, and *vice versa*.

**IMPACT STATEMENT:** Adaptation to changing environments requires a source of genetic variation. When environments change quickly, species often rely on variation that is already present – so-called standing genetic variation – because new adaptive mutations are just too rare. The threespine stickleback, a small fish species living throughout the Northern Hemisphere, is well-known for its ability to rapidly adapt to new environments. Populations living in coastal oceans are heavily armored with bony plates and spines that protect them from predators. These marine populations have repeatedly invaded and adapted to freshwater environments, losing much of their armor and changing in shape, size, color, and behavior.

Adaptation to freshwater environments can occur in mere decades and probably involves lots of standing genetic variation. Indeed, one of the clearest examples we have of adaptation from standing genetic variation comes from a gene, *eda*, that controls the shifts in armor plating. This discovery involved two surprises that continue to shape our understanding of the genetics of adaptation. First, freshwater stickleback from across the Northern Hemisphere share the same version, or allele, of this gene. Second, the ‘marine’ and ‘freshwater’ alleles arose millions of years ago, even though the freshwater populations studied arose much more recently. While it has been hypothesized that other genes in the stickleback genome may share these patterns, large-scale surveys of genomic variation have been unable to test this prediction directly.

Here, we use new sequencing technologies to survey DNA sequence variation across the stickleback genome for patterns like those at the *eda* gene. We find that *every* region of the genome associated with marine-freshwater genetic differences shares this pattern to some degree. Moreover, many of these regions are as old or older than *eda*, stretching back over 10 million years in the past and perhaps even predating the species we now call the threespine stickleback. We conclude that natural selection has maintained this variation over geological timescales and that the same alleles we observe in freshwater stickleback today are the same as those under selection in ancient, now-extinct freshwater habitats. Our findings highlight the need to understand evolution on macroevolutionary timescales to understand and predict adaptation happening in the present day.

## INTRODUCTION

The mode and tempo of adaptive evolution depend on the sources of genetic variation affecting fitness (Wright 1932; Orr 2005). While new mutation is the ultimate origin of all genetic variation, recent studies of adaptation in the wild have documented adaptive genetic variation that was either segregating in the ancestral population as standing genetic variation (SGV)(Barrett & Schluter 2008; Domingues *et al.* 2012; Schrider & Kern 2017), or introgressed from a separate population or species (Huerta-Sánchez *et al.* 2014; Fontaine *et al.* 2015). The use of SGV during evolution appears particularly important when dramatic responses to selection occur on ecological timescales, in dozens of generations or fewer (Barrett & Schluter 2008). When environments change rapidly, SGV can propel rapid evolution in ecologically relevant traits even in populations of long-lived organisms like Darwin's finches (Grant & Grant 2002), monkeyflowers (Wright *et al.* 2013), and threespine stickleback fish (Colosimo *et al.* 2005).

The contribution of SGV to rapid divergence has important consequences for our understanding of evolutionary genetics. Existing genetic variants have evolutionary histories that are often unknown, but which may none-the-less have significant impacts on subsequent adaptation (Kirkpatrick & Barton 2006; Wright *et al.* 2013). The abundance, genomic distribution, and fitness effects (Charlesworth *et al.* 1993; Colosimo *et al.* 2005; Kirkpatrick & Barton 2006; Linnen *et al.* 2009; Stankowski & Streisfeld 2015) of SGV are themselves the products of evolution, and their unknown history raises fascinating questions for the genetics of adaptation in the wild. When did adaptive variants originally arise? How are they structured, across both geography and the genome? Which evolutionary forces shaped their current distribution and how might this evolutionary history channel future evolutionary change?

Answers to these questions are critical for our understanding of the importance of SGV in nature, as well as our ability to predict the paths available to adaptation on ecological timescales (Wright *et al.* 2013). Biologists are beginning to probe evolutionary histories of SGV using genome-wide sequence variation across multiple individuals in numerous populations (Pease *et al.* 2016), but this level of inference has been unavailable for most natural systems because of methodological limitations that remove phase information (e.g. pool-seq: Schlotterer *et al.* 2014) or produce very short reads (e.g. RAD-seq: Davey *et al.* 2011). Here, we investigate the structure and evolutionary history of divergent SGV by modifying the original sheared RAD-seq method to generate ~700 bp haplotypes at tens of thousands of loci sampled across the stickleback genome. This approach allows us to accurately measure sequence variation and estimate divergence times across the genome. By collecting more detailed sequence information at each RAD locus, this approach also provides more accurate estimates of polymorphism and divergence at each locus, and with far smaller sample sizes, compared to traditional short-read methods (Nei 1987 chapters 10 and 13; Wakeley 2009; Cruickshank & Hahn 2014).

SGV has long been postulated to be critical to adaptation in stickleback, and several recent population genomic studies have supported this hypothesis (Hohenlohe *et al.* 2010; Jones *et al.* 2012; Roesti *et al.* 2015; Samuk *et al.* 2017). Marine stickleback have repeatedly colonized freshwater lakes and streams (Bell & Foster 1994b; Jones *et al.* 2012; Wund *et al.* 2016), and adaptive divergence in isolated freshwater habitats is highly parallel at the phenotypic (Colosimo *et al.* 2004; Cresko *et al.* 2004a) and genomic levels (Hohenlohe *et al.* 2010; Jones *et al.* 2012; but see Stuart *et al.* 2017). In addition, analyses of haplotype variation at the genes *eda* (Colosimo *et al.* 2005; Roesti *et al.* 2014) and *atp1a1* (Roesti *et al.* 2014) present two clear results: separate freshwater populations share common ‘freshwater’ haplotypes that are identical-by-descent (IBD), and sequence divergence between the major marine and freshwater haplogroups suggests their ancient origins – perhaps over two million years ago in the case of *eda* (Colosimo *et al.* 2005). While intriguing, it is not clear whether the deep evolutionary histories of these loci are outliers or representative of more widespread ancient history across the genome. Furthermore, although recent population genomic studies have made important contributions to identifying that SGV across the genome, the short reads employed limit the accuracy of genealogical inference across the genome.

To address fundamental questions of genealogical relationships and molecular evolution in stickleback, we utilize the new RAD-seq haplotyping approach to assay genome-wide variation associated with adaptive divergence in two young freshwater ponds, which formed during the end-Pleistocene glacial retreat (c. 12,000 years ago: Francis *et al.* 1986; Cresko *et al.* 2004a, Fig. 1). In addition, we generated a *de novo* genome assembly of the sister taxon ninespine stickleback (*Pungitius pungitius*), allowing us to estimate divergence times for genealogies across the genome. Our results clearly demonstrate that the previous findings of deep evolutionary history based upon candidate loci are not unique but in fact the rule. A suite of adaptive variation structured into distinct marine and freshwater haplotypes that evolved over millions of years forms the foundation of a deep pool of SGV that undergirds repeated and rapid evolution in stickleback.

**Figure 1.**
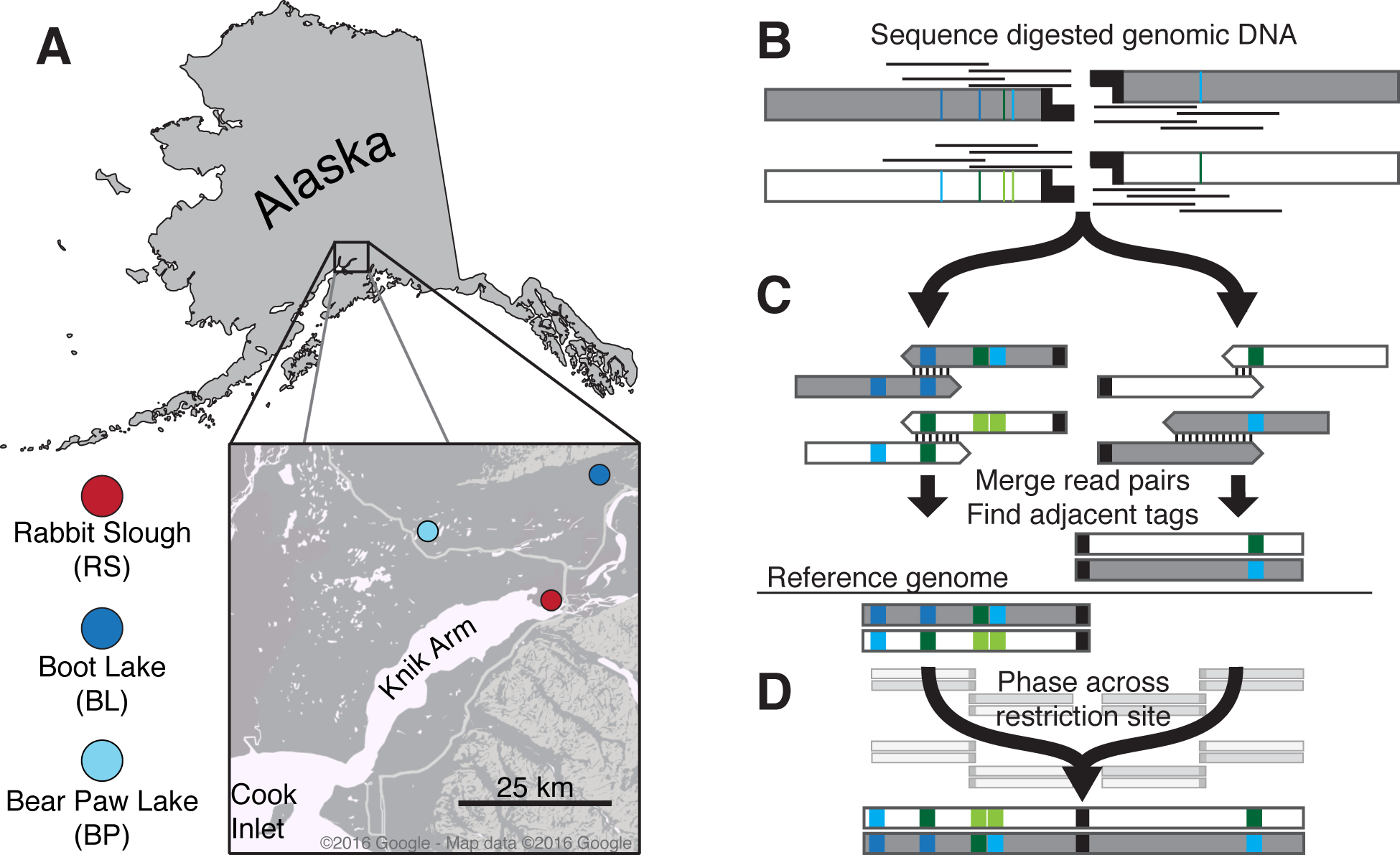
Stickleback sampling and RAD sequencing to measure haplotype variation. A) Threespine stickleback sampling locations in this study. Colors represent habitat type: red: marine; blue: freshwater. B-D: We modified the original RAD-seq protocol to generate local haplotypes. Colored bars represent polymorphic sites. For a detailed description of haplotype construction, see *Methods*. B) Overlapping paired-end reads are anchored to *PstI* restriction sites. C) Paired reads mapping to each halfsite are merged into contigs. Contigs mapping to the same restriction site are identified by alignment to the reference genome. D) Sequences from each half of a restriction site are phased to generate a single RAD locus. RAD tags in the background represent multiple genotypes used in phasing.

## METHODS

### Sample collection

Wild threespine stickleback were collected from Rabbit Slough (N 61.5595, W 149.2583), Boot Lake (N 61.7167, W 149.1167), and Bear Paw Lake (N 61.6139, W 149.7539). Rabbit Slough is an offshoot of the Knik Arm of Cook Inlet and is known to be populated by anadromous populations of stickleback that are stereotypically oceanic in phenotype and genotype (Cresko *et al.* 2004b; Hohenlohe *et al.* 2010). Boot Lake and Bear Paw Lake are both shallow lakes formed during the end-Pleistocene glacial retreat. Fish were collected in the summers of 2009 (Rabbit Slough), 2010 (Bear Paw Lake), and 2014 (Boot Lake) using wire minnow traps and euthanized *in situ* with Tricaine solution. Euthanized fish were immediately fixed in 95% ethanol and shipped to the Cresko Laboratory at the University of Oregon (Eugene, OR, USA). DNA was extracted from fin clips preserved in 95% ethanol using either Qiagen DNeasy spin column extraction kits or Ampure magnetic beads (Beckman Coulter, Inc) following manufacturer's instructions. Yields averaged 1-2 μg DNA per extraction (~30 mg tissue). Treatment of animals followed protocols approved the University of Oregon Institutional Animal Care and Use Committee (IACUC).

### Sequencing strategy and rationale

We designed our sequencing to maximize detection of sequence variation and divergence, with the ultimate goal being the estimation of absolute divergence times of marine and freshwater haplogroups. Previous work by us and others using short sequence reads provided clear evidence of changes in relative frequencies of alleles across stickleback populations (Hohenlohe *et al.* 2010; Roesti *et al.* 2014; Lescak *et al.* 2015; Roesti *et al.* 2015), but could not sufficiently address questions of haplotype ages. We therefore designed a RAD sequencing approach to (1) accurately estimate sequence diversity within and divergence between threespine stickleback ecotypes and (2) recover sufficient RAD loci that map unambiguously to an outgroup genome sequence from the ninespine stickleback that we could confidently compare diversity within threespine stickleback to divergence from the ninespine stickleback.

To achieve our aims, we designed a sequencing method to produce phased haplotypes of ~700 bp at each RAD locus (Fig. 1B-D) and to sample the genome densely enough to identify signatures of selection after the likely dropout of RAD loci without clear homology in the ninespine stickleback genome. We used the single-digest, sheared RAD approach to limit biases in our estimates of sequence diversity. RAD-seq has known biases due to mutations in restriction sites causing allele dropout (Arnold *et al.* 2013; Gautier *et al.* 2013), the potential for which increases with increasing sequence divergence and leads to underestimates of genetic diversity. Diversity estimates are, however, substantially more accurate with sheared RAD-seq compared to other RAD-seq approaches (e.g. double-digest RAD-seq: Peterson *et al.* 2012). Importantly for the coalescent analyses we present here, such allele dropout is unlikely to affect estimates of overall divergence across the clade of alleles. When in the rare cases it does, the bias is toward *underestimation* of the divergence age (Arnold *et al.* 2013), which would make our findings of deep divergence even more striking.

Our sequencing design facilitated accurate inference of sequence variation even with smaller population samples than are typical among population genomic studies. While allele frequency-based statistics like F_ST_ have particularly high variance with small sample sizes (Willing *et al.* 2012), our study is fortunate to be built upon numerous properly powered, previous population genomic studies in stickleback including in these populations. The genome-wide patterns of F_ST_ we observed using our new approach closely matched multiple previous studies (Hohenlohe *et al.* 2010; Jones *et al.* 2012, Fig. 2A). Because of this extensive body of previous work, we relied on F_ST_ only to draw inference of larger genomic regions containing tens or hundreds of RAD loci. Instead, as stated above the focus of this work is to extend these previous findings by addressing the ages of allelic divergence. We therefore do not expect the higher variance associated with smaller sample size to qualitatively influence our results. Importantly, estimation of sequence diversity (*π*) (Nei 1987) and divergence (*d*_*XY*_) (Nei 1987; Cruickshank & Hahn 2014) at a given locus improves greatly with increases in sequence length. Using equations 10.9 and 13.83 from Nei (1987; Box 1 in Cruickshank & Hahn 2014), the predicted sampling variances in both *π* and *d*_*XY*_ using 700 bp sequences in five individuals are lower than those obtained using standard 100 bp sequences at any sample size (Suppl. Fig. S1). Therefore, not only is this novel application of RAD-seq ideally suited for our questions, our findings show that this approach may significantly decrease the necessary sample size, and thus resource expenditure, for many population genomic studies.

**Figure 2.**
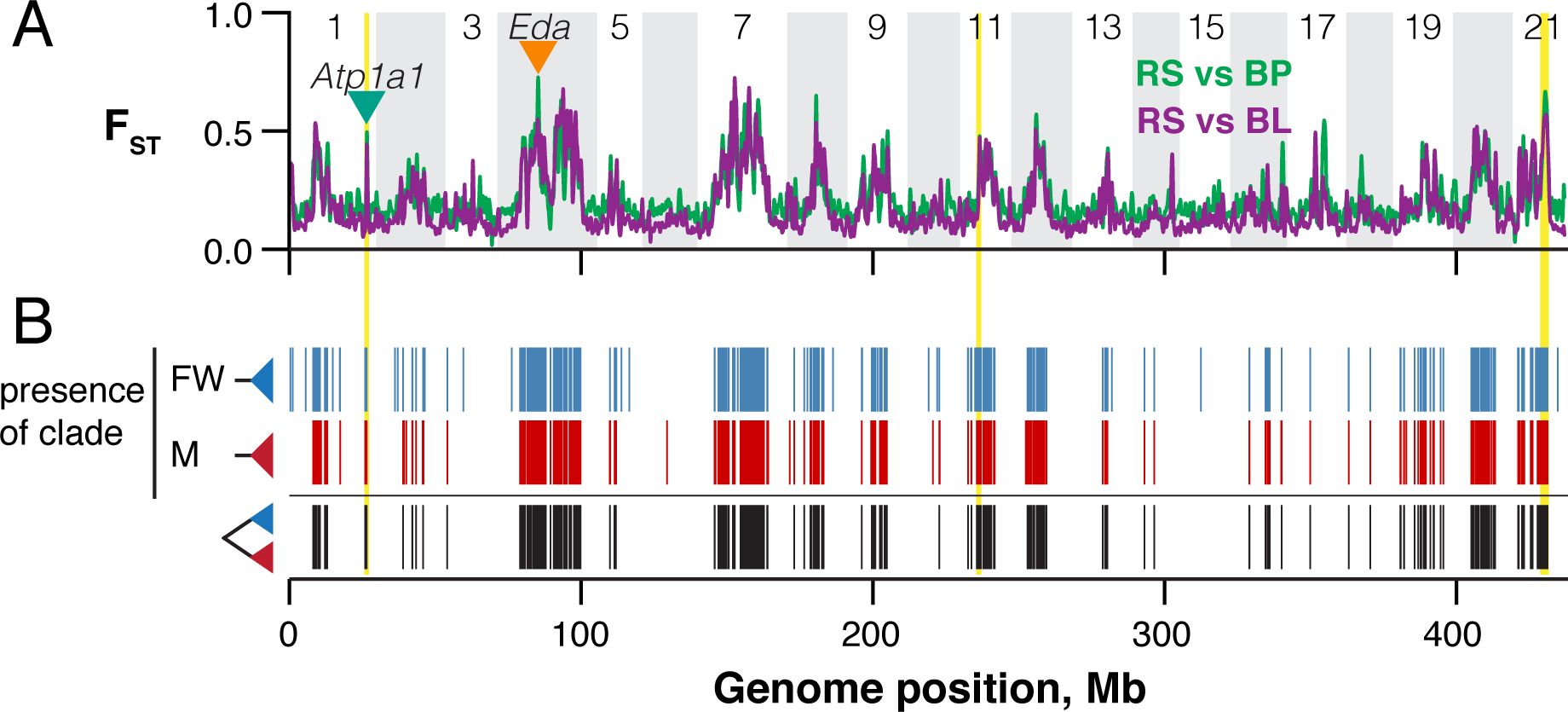
The genealogical structure of parallel genomic divergence. A) Genome-wide F_ST_ for both marine-freshwater comparisons was kernel-smoothed using a normally distributed kernel with a window size of 500 kb. Inverted triangles indicate the locations of two genes known to show extensive marine-freshwater haplotype divergence, *Eda* and *Atp1a1*. Three chromosomal inversions are highlighted in yellow. B) Lineage sorting patterns were identified from maximum clade credibility trees for each RAD locus. Blue bars: haplotypes from both freshwater populations form a single monophyletic group; red: haplotypes from the marine population form a monophyletic group; black: A RAD locus is structured into reciprocally monophyletic marine and freshwater haplogroups.

### Library preparation

To identify sufficient sequence variation at a RAD locus, and to simplify downstream sequence processing and analysis, we took advantage of longer sequencing reads available on newer Illumina platforms and the phase information captured by paired-end sequencing. We generated RAD libraries from these samples using the single-digest sheared RAD protocol from Baird et al. (2008) with the following specifications and adjustments: 1 μg of genomic DNA per fish was digested with the restriction enzyme *PstI-HF* (New England Biolabs), followed by ligation to P1 Illumina adaptors with 6 bp inline barcodes. Ligated samples were multiplexed and sheared by sonication in a Bioruptor (Diagenode). To ensure that most of our paired-end reads would overlap unambiguously and produce longer contiguous sequences, we selected a narrow fragment size range of 425-475 bp. The remainder of the protocol was per Baird et al. (Baird *et al.* 2008b). All fish were sequenced on an Illumina HiSeq 2500 using paired-end 250 bp sequencing reads at the University of Oregon's Genomics and Cell Characterization Core Facility (GC3F).

### Sequence processing

Raw Illumina sequence reads were demultiplexed, cleaned, and processed primarily using the Stacks v1.46 pipeline (Catchen *et al.* 2011; Catchen *et al.* 2013a). Paired-end reads were demultiplexed with `process_shortreads` and cleaned using `process_radtags` using default criteria (throughout this document, names of scripts, programs, functions, and command-line arguments will appear in `fixed-width font`). Overlapping read pairs were then merged with `fastq-join` (Aronesty 2011). Pairs that failed to merge were removed from further analysis. To retain the majority of the sequence data for analysis in Stacks and still maintain adequate contig lengths, merged contigs were trimmed to 350 bp and all contigs shorter than 350 bp were discarded. We aligned these contigs to the stickleback reference genome (Jones *et al.* 2012; Glazer *et al.* 2015) using `bbmap` v35.69 with the most sensitive alignment settings (‘`vslow=t`’; http://jgi.doe.gov/data-and-tools/bbtools/) and required that contigs mapped uniquely to the reference. We then used the `pstacks`, `cstacks`, and `sstacks` components of the Stacks pipeline to identify RAD-tags and call SNPs using the maximum likelihood algorithm implemented in `pstacks`, create a catalog of RAD tags across individuals, and match tags across individuals. All data were then passed through the Stacks error correction module `rxstacks` to prune unlikely haplotypes. We ran the Stacks component program `populations` on the final dataset to filter loci genotyped in fewer than four individuals in each population and to create output files for sequence analysis. We use the naming conventions of Baird et al. (2008a): A “RAD tag” refers to sequence generated from a single end of a restriction site and the pair of RAD tags sequenced at a restriction site comprises a “RAD locus” (Fig. 1D).

We used the program `phase` v2.1 (Stephens *et al.* 2001; Stephens & Scheet 2005) to phase pairs of RAD tags originating from the same restriction site. We coded haplotypes present at each RAD tag, which often contain multiple SNPs, into multiallelic genotypes. This both simplified and reduced computing time for the phasing process. We also performed coalescent simulations to generate, ‘cut’, and re-phase haplotypes to demonstrate the high accuracy of this method using sequences and sample sizes similar to those in this study (Suppl. Fig. S2). Custom Python scripts automated this process and are included as supplementary files. We required that each individual had at least one sequenced haplotype at each tag for phasing to be attempted. If a sample had called genotypes at only one tag in the pair, the sample was removed from further analysis of that locus. The resultant phased haplotypes were used to generate sequence alignments for import into BEAST.

We recovered a total of 236,787 RAD tags after filtering, mapping to 151,813 *PstI* restriction sites. At 84,974 restriction sites, we recovered and successfully phased adjacent RAD tags (169,948 RAD tags) into single RAD loci. RAD tags with no variable sites were simply concatenated to the adjacent tag to form a single locus. We retained these 84,974 RAD loci for our analysis. For population genetic analyses, inclusion of singleton (i.e. unpaired) RAD tags did not qualitatively change our results. We chose to restrict genealogical analyses to loci of uniform length and to use the same set of loci in analyses of polymorphism and gene tree topologies.

### Ninespine stickleback genome assembly

In order to estimate the T_MRCA_ of threespine stickleback RAD alleles, we used the ninespine stickleback (*Pungitius pungitius*) as an outgroup. RAD sequence analysis, however, relies on the presence of homologous restriction sites among sampled individuals and results in null alleles when mutations occur within a restriction site(Arnold *et al.* 2013). Because this probability increases with greater evolutionary distance among sampled sequences, we elected to use RAD-seq to only estimate sequence variation within the threespine stickleback. We then generated a contig-level *de novo* ninespine stickleback genome assembly from a single ninespine stickleback individual from St. Lawrence Island, Alaska (collected by J. Postlethwait) using DISCOVAR *de novo* revision 52488 (https://software.broadinstitute.org/software/discovar/blog/). We used this single ninespine stickleback haplotype to estimate threespine-ninespine sequence divergence and time calibrate coalescence times within the threespine stickleback. DISCOVAR de novo requires a single shotgun library of paired-end 250-bp sequence reads from short-insert-length DNA fragments. High molecular weight genomic DNA was extracted from an ethanol-preserved fin clip by proteinase K digestion followed by DNA extraction with Ampure magnetic beads. Purified genomic DNA was mechanically sheared by sonication and size selected to a range of 200-800 bp by gel electrophoresis and extraction. We selected this fragment range to agree with the recommendations for *de novo* assembly using DISCOVAR *de novo*. This library was sequenced on a single lane of an Illumina HiSeq2500 at the University of Oregon's Genomics and Cell Characterization Core Facility (GC3F: https://gc3f.uoregon.edu/). We assembled the draft ninespine stickleback genome using DISCOVAR de novo. Raw sequence read pairs were first quality filtered and adaptor sequence contamination removed using the program `process_shortreads`, which is included in the Stacks analysis pipeline (Catchen *et al.* 2013b). We ran the genome assembly on the University of Oregon's Applied Computational Instrument for Scientific Synthesis (ACISS: http://acisscomputing.uoregon.edu).

### Alignment of RAD tags to the ninespine assembly

We included the single ninespine stickleback haplotype into our sequence analyses by aligning a single phased threespine stickleback RAD haplotype from each locus to the ninespine genome assembly. For those that aligned uniquely (59,254 RAD loci), we used a custom Python script to parse the alignment fields of the output BAM file (Li *et al.* 2009) and reconstruct the ninespine haplotype by introducing threespine-ninespine substitutions into the threespine RAD locus sequence. The final dataset consists of 57,992 RAD loci that mapped to the 21 threespine stickleback chromosomes and aligned uniquely to the ninespine assembly.

### Lineage sorting and time to the most recent common ancestor

Allelic divergence can occur by multiple modes of lineage sorting during adaptation. To identify patterns of lineage sorting associated with freshwater colonization, we analyzed gene tree topologies at all RAD loci using BEAST v. 1.7 (Drummond & Rambaut 2007; Drummond *et al.* 2012). We chose BEAST because it co-estimates tree topologies and node ages for sequenced genomic loci. BEAST does not explicitly perform model selection, and this may affect divergence time estimates in genomic regions under direct or indirect selection. However, other methods developed to estimate the age of adaptive alleles model evolutionary scenarios that are likely not relevant to the evolutionary histories we infer here. First, some models assume a recent origin of an adaptive allele compared to adjacent genomic variation (Peter *et al.* 2012; Ormond *et al.* 2016), which is the opposite of what we describe here, so that measures of variation at linked sites and the decay of linkage disequilibrium can be used to estimate when a sweep began. Selection in the stickleback populations we study likely acted on SGV, as has been supported by previous studies, and we hypothesize that this SGV may be quite old. Therefore, adaptive alleles already existed on distinct haplotype backgrounds, which masks the differences between selected and linked neutral sites.

Second, a recent model developed to infer ages of standing genetic variants assumes that the variant was evolving neutrally at some point during its trajectory through a population (Peter *et al.* 2012). This assumption is unlikely for many of the loci we detect here, except in the very distant past and for those loci that have evolved recently arose in genomic regions already heavily influenced by selection. Rather, the patterns of haplotype variation we observed in the genomic regions that differentiate marine and freshwater populations reflect long-term maintenance and isolation of separate haplogroups that mimics population structure and even speciation, with selective sweeps being important but constituting a small minority of the time these haplotypes have segregating in the stickleback metapopulation. For all of these reasons we therefore chose to estimate tree topologies and divergence times with BEAST, which makes minimal assumptions regarding specific evolutionary processes.

We used blanket parameters and priors for BEAST analyses across all RAD loci. Markov chain Monte Carlo (MCMC) runs of 1,000,000 states were specified, and trees logged every 100 states. We used a coalescent tree prior and the GTR+Γ substitution model with four rate categories and uniform priors for all substitution rates. We identified evidence of lineage sorting by using the program `treeannotator` v1.7.5 to select the maximum clade credibility (MCC) tree for each RAD locus and the `is.monophyletic()` function included in the R package ‘ape’ v3.0 (Paradis *et al.* 2004; Popescu *et al.* 2012). We determined for each MCC tree whether tips originating from marine (RS) or freshwater (BL+BP) formed monophyletic clades.

To convert node ages estimated in BEAST into divergence times, in years, we assumed a 15 million-year divergence time between threespine and ninespine stickleback at each RAD locus (Aldenhoven *et al.* 2010). The T_MRCA_ of all alleles in each gene tree was set at 15 Mya and each node age of interest was converted into years relative to the total height of the tree. Additionally, to use the ninespine stickleback as an outgroup, we required that threespine stickleback haplotypes at a RAD locus were monophyletic to the exclusion of the ninespine haplotype. Doing so reduced our analysis to 49,672 RAD loci for analyses included in Fig. 4 of the main text. RAD loci not showing this pattern of lineage sorting did not show evidence of a genome-wide correlation with marine-freshwater divergence and thus do not impact the assertions in the main text. We used medians of the posterior distributions as point estimates of T_MRCA_ for each RAD locus. Because of the somewhat limited information from any single RAD locus, and because the facts of the genealogical process mean that the true T_MRCA_ at any locus likely differs from the 15 My estimate (Kingman 1982a, b; Tajima 1983), we do not rely heavily on T_MRCA_ estimates at individual RAD loci. Rather, we use these estimates to understand patterns of broad patterns of ancestry throughout the threespine stickleback genome — spatially along chromosomes and genome-wide patterns.

We determined T_MRCA_ outlier genomic regions by permuting and kernel smoothing the genomic distribution of T_MRCA_ estimates using the same window sizes as we present in the main text. Windows where the actual T_MRCA_ exceeded 99.9% of permuted windows were considered outliers. This method controls for the local density of RAD loci (poorly sampled regions will have larger confidence bands) and the size of the windows used.

### Sequence diversity and haplotype networks

We quantified sequence diversity within and among populations and sequence divergence between populations using R v3 (R Core Team 2016). We used the R package ‘ape’ (Paradis *et al.* 2004) to compute pairwise distance matrices for all alleles at each RAD locus and used these matrices to calculate the average pairwise nucleotide distances, *π*, within and among populations along with *d*_*XY*_, the average pairwise distance between two sequences using only across-population comparisons (Nei 1987). We also calculated the haplotype-based F_ST_ from Hudson et al. (1992) implemented in the R package ‘PopGenome’ v2.2.4 (Pfeifer *et al.* 2014). We used permutation tests written in R to identify differences in variation within- and between-habitat type at divergent RAD loci versus the genome-wide distributions. Mann-Whitney-Wilcoxon tests implemented in R were used to identify variation in genome-wide diversity among populations and habitat types.

We constructed haplotype networks of the RAD loci at *eda* and *atp1a1* using the infinite sites model with the function `haploNet()` in the R package ‘pegas’ (Paradis 2010). The *atp1a1* network was constructed from from a RAD locus spanning exon 15 of *atp1a1* and including portions of introns 14 and 15 at (chr1:21,726,729-21,727,381 [BROAD S1, v89]; chr1: 26,258,117-26,257,465 [re-scaffolding from Glazer, *et al* (2015)]). The *eda* network spans exon 2 and portions of introns 1 and 3 of *eda* (chr4: 12,808,396-12,809,030).

### Code availability

Scripts used to phase RAD-tags, summarize gene trees, calculate population genetic statistics, and produce figures and statistics presented in paper are available at https://github.com/thomnelson/ancient-divergence. Scripts for processing raw sequence data are available from the authors upon request.

### Data availability

Raw sequence data supporting these findings are available on the Sequence Read Archive at PRJNAXXXXXX. The final datasets needed to reproduce the figures and statistics presented in the paper are available at https://github.com/thomnelson/ancientdivergence.

## RESULTS AND DISCUSSION

Parallel adaptation to freshwater environments has been a major theme of stickleback evolutionary history (Bell & Foster 1994a). Stereotypical morphological changes to, for example, bony armor(Colosimo *et al.* 2004) and craniofacial structures (Kimmel *et al.* 2005) presumably reflect adaptation to similar selective regimes (Reimchen 1994; Arnegard *et al.* 2014). These phenotypic changes are accompanied by parallel genomic divergence (Hohenlohe *et al.* 2010; Jones *et al.* 2012), which involves large regions spanning many megabases (Schluter & Conte 2009; Roesti *et al.* 2014), including multiple chromosomal inversions (Jones *et al.* 2012). The leading hypothesis for the genetics of parallel divergence in stickleback posits that distinct freshwater-adaptive haplotypes that are identical-by-descent (IBD) are shared among freshwater populations due to historical gene flow between marine and freshwater populations (Schluter & Conte 2009). We tested for the presence of these haplotypes directly and at a genomic scale.

### Parallel divergence involves a shared suite of haplotypes genome-wide

Our sequencing strategy produced 57,992 RAD loci, with 690 potential variable sites each, present across the three threespine stickleback populations and aligned to the ninespine stickleback genome assembly. These data comprise over 40 Mb of sequence, or nearly 10% of the threespine stickleback genome (9.5% of 419 Mb assigned to chromosomes) (Jones *et al.* 2012; Glazer *et al.* 2015). All loci we recovered were polymorphic and we observed a median of seven segregating sites per locus (range: 2-155, Suppl. Fig. S3, Suppl. Table 1). By including haplotypes from all three populations in these genealogical analyses, we were able to jointly calculate population genetic statistics (F_ST_, *π*, *d*_*XY*_) and identify patterns of identity-by-descent (IBD) among populations, which we defined as haplotypes from two populations forming a monophyletic group to the exclusion of the third population.

We find that parallel population genomic divergence in the two freshwater pond populations consistently involved haplotypes that were identical-by-descent (IBD) among both freshwater populations (Fig. 2). Background F_ST_ between populations ranged from 0.139-0.226, with genome-wide differentiation between the freshwater populations BL and BP being highest (F_ST(RS-BL)_ = 0.139, F_ST(RS-BP)_ = 0.194, F_ST(BL-BP)_ = 0.226; two-sided Mann-Whitney test for all pairwise comparisons: *p* ≤ 1×10^−10^). The degree and genomic distribution of pairwise F_ST_ between the BL, BP, and RS populations were similar to those previously reported (Hohenlohe *et al.* 2010). This similarity included marine-freshwater F_ST_ outlier regions on chromosome 4 over a broad span in which the *eda* gene is embedded (orange triangle in Fig. 2A), and three regions now known to be associated with chromosomal inversions on chromosomes 1, 11, and 21 (yellow bars in Fig. 2; hereafter referred to as *inv1*, *inv11*, and *inv21*). The gene *atp1a1* (green triangle in Fig. 2A) is contained within *inv1*. As expected, we found distinct haplogroups associated with marine and freshwater habitats at both *eda* and *atp1a1* (Fig. 3, insets).

**Figure 3.**
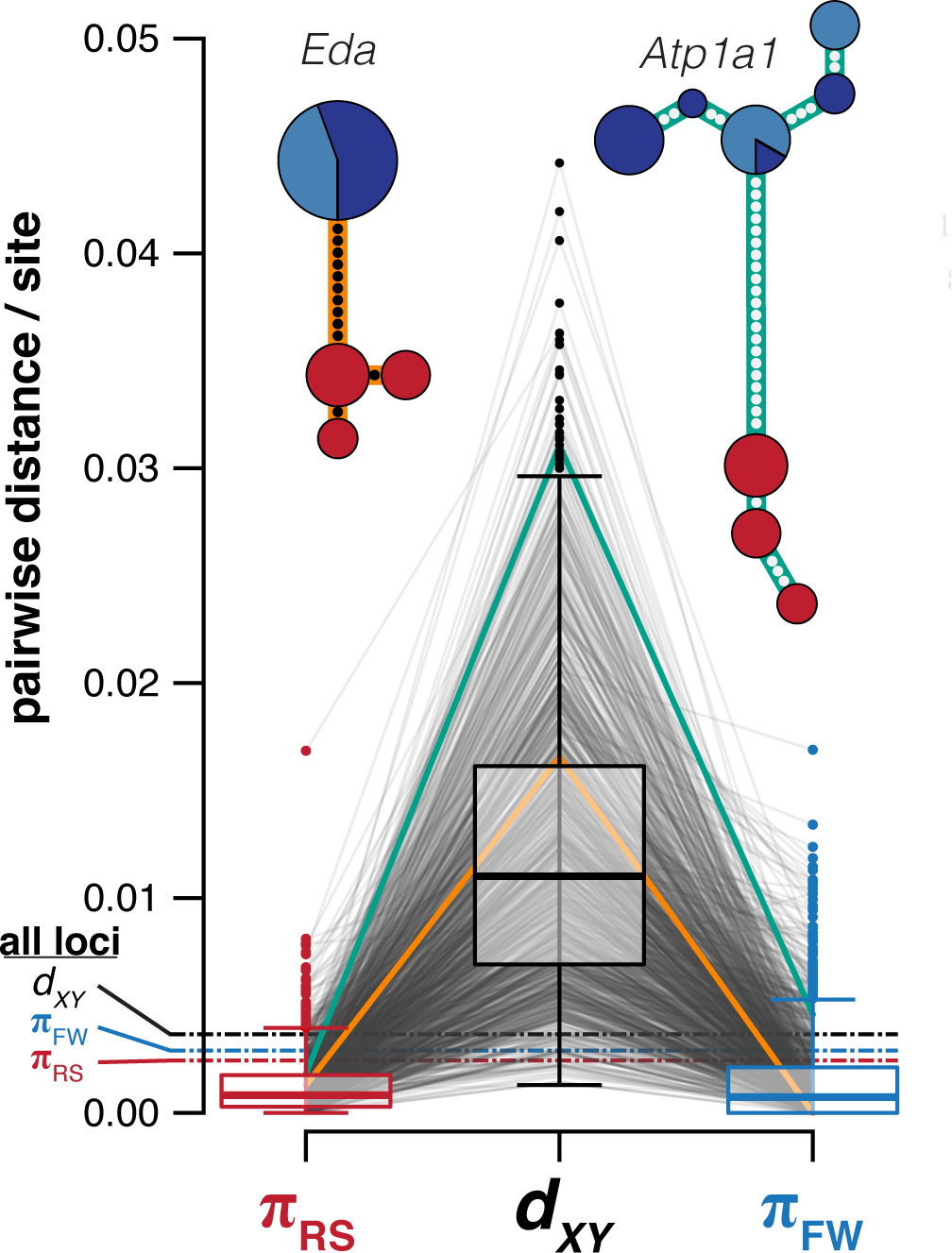
Extensive sequence divergence between marine and freshwater haplogroups accompanies reciprocal monophyly. For each reciprocally monophyletic RAD locus, we calculated sequence variation (*π*) within and sequence divergence between habitat types (*d*_*XY*_). Each RAD locus is shown as a pair of lines connecting estimates of π and *d*_*XY*_. Boxplots show distributions across all reciprocally monophyletic RAD loci: Boxes are upper and lower quartiles, including the median; whiskers extend to 1.5x interquartile range. Dashed lines are the genome-wide medians. Single RAD loci from within the transcribed regions of *Eda* and *Atp1a1* are shown as gold and green lines, respectively, and presented as haplotype networks. Dots represent mutational steps. Circle sizes indicate the number of haplotypes and colors indicate population of origin as in Figure 1. Each network = 29 haplotypes.

Strikingly, this finding of habitat specific haplogroups was not at all unique to these well studied genes or chromosomal inversions. The two isolated freshwater populations shared IBD haplotypes within all common marine-freshwater F_ST_ peaks even though IBD was rare elsewhere (Fig. 2B). Furthermore, we observed a separate clade of haplotypes representing the marine RS population at the majority (1129 of 2172, 52%) of RAD loci showing freshwater IBD. The result was a genome-wide pattern of reciprocal monophyly between marine and freshwater haplotypes. Notably, this is the same genealogical structure previously reported at *eda* (Colosimo *et al.* 2005; Roesti *et al.* 2014) and *atp1a1* (Roesti *et al.* 2014), demonstrating that these loci are but a small part of a genome-wide suite of genetic variation sharing similar habitat-specific evolutionary histories, and the previous documentation of their genealogies was a harbinger of a much more extensive pattern across the genome revealed here. Hereafter, we refer collectively to this class of RAD loci as ‘divergent loci’.

### Adaptive marine-freshwater sequence divergence involves ancient allelic origins

Because the genealogical structure of divergence across the genome mirrors that at *eda* and *atp1a1*, we asked whether levels of sequence variation and divergence also showed consistent genomic patterns. At all RAD loci we therefore calculated *π* within each population, as well as in the combined freshwater populations, and *d*_*XY*_ between marine and freshwater habitat types. Genome-wide diversity was similar across populations and habitat types (mean *π*_RS_ = 0.0032, *π*_BL_ = 0.0034, *π*_BP_ = 0.0026, *π*_FW_ = 0.0038) and comparable to previous estimates (Hohenlohe *et al.* 2010). Likewise, genome-wide *d*_*XY*_ among habitat types was modest (0.0049) when compared to *π* across all populations (*π* = 0.0042, two-sided Mann-Whitney test: *p* ≤ 1×10^−10^; Suppl. Fig. S4). Among divergent loci, however, we observed reductions in diversity in both habitats (mean *π*_RS-divergent_ = 0.0012, *π*_RS-divergent_ = 0.0016, two-sided permutation test: *p* ≤ 1×10^−4^, Fig. 3), indicating natural selection in both habitats. Sequence divergence associated with reciprocal monophyly was striking, however, averaging nearly three times the genome-wide mean (mean *d*_*XY*-divergent_ = 0.0124). This divergence ranged more than an order of magnitude (0.0013–0.0442), from substantially lower than the genome-wide average to ten times greater than the average. These findings indicate that much of the genetic variation underlying adaptive divergence is vastly older than the diverging freshwater populations themselves. Not only was adaptive variation standing and structured by habitat, but it has been segregating and accumulating for millennia.

These data clearly support the hypothesis of Schluter and Conte (2009) of ancient haplotypes ‘transported’ among freshwater populations. Much of the divergence we observed was ancient in origin, with levels of sequence divergence at some RAD loci exceeding that observed at *eda* (Fig. 3, gold line) and suggestive of divergence times of at least two million years ago (Colosimo *et al.* 2005). Our observation that sequence variation was consistently reduced in both habitat types emphasizes that alternative haplotypes at these loci are likely selected for in the marine population as well as the freshwater. These alternative fitness optima — driven by divergent ecologies — provide a favorable landscape for the maintenance of variation (Charlesworth *et al.* 1997; Lenormand 2002), but also lead to a more potent barrier to gene flow among freshwater populations if there are fitness consequences in the marine habitat for stickleback carrying freshwater-adaptive variation. Conditional fitness effects through genetic interactions (i.e. dominance or epistasis: Otto & Bourguet 1999; Phillips 2008) and genotype-by-environment interactions (McGuigan *et al.* 2011) could potentially extend the residence time of freshwater haplotypes in the marine habitat. Future work should consider the phenotypic effects of divergently adaptive variation in different external environments (McGuigan *et al.* 2011; McCairns & Bernatchez 2012).

Adaptive divergence between marine and freshwater stickleback genomes is likely ongoing, with recently derived alleles arising on already highly divergent genomic backgrounds. We found reciprocal monophyly associated with a spectrum of sequence divergence, including a substantial fraction of divergent loci (11.0%, 124/1129) with *d*_*XY*_ below the genome-wide average. Thus, ongoing marine-freshwater ecological divergence may continue to yield additional marine-freshwater genomic divergence. Moreover, while this younger variation is shared between the freshwater populations in this study, and localizes to genomic regions of divergence shared globally (Jones *et al.* 2012), some adaptive variants may be distributed only locally (e.g. limited to southern Alaska or the eastern Pacific basin). Global surveys of shared variation have been performed (Jones *et al.* 2012), but future work in this system should quantify the distributions of locally or regionally limited genomic variation involved in ecological divergence, because regional pools of variation may contribute substantially to stickleback genomic and phenotypic diversity (Stuart *et al.* 2017).

### Habitat associated genomic divergence is as old as the threespine stickleback species

Sequence divergence provides an important relative, but ultimately incomplete, evolutionary timescale. To more directly compare the timescales of ecological adaptation and genomic evolution, we translated patterns of sequence variation into the time to the most recent common ancestor (T_MRCA_) of allelic variation, in years. To do so, we performed a *de novo* genome assembly of the ninespine stickleback (*Pungitius pungitius*), a member of the Gasterosteidae that diverged from the threespine stickleback lineage approximately 15 million years ago (MYA) (Aldenhoven *et al.* 2010) (Fig. 4A, Suppl. Table 2). We then aligned our RAD dataset to this assembly and estimated gene trees for each alignment with BEAST (Drummond *et al.* 2012), setting divergence to the ninespine stickleback at 15 MYA (see Methods).

**Figure 4.**
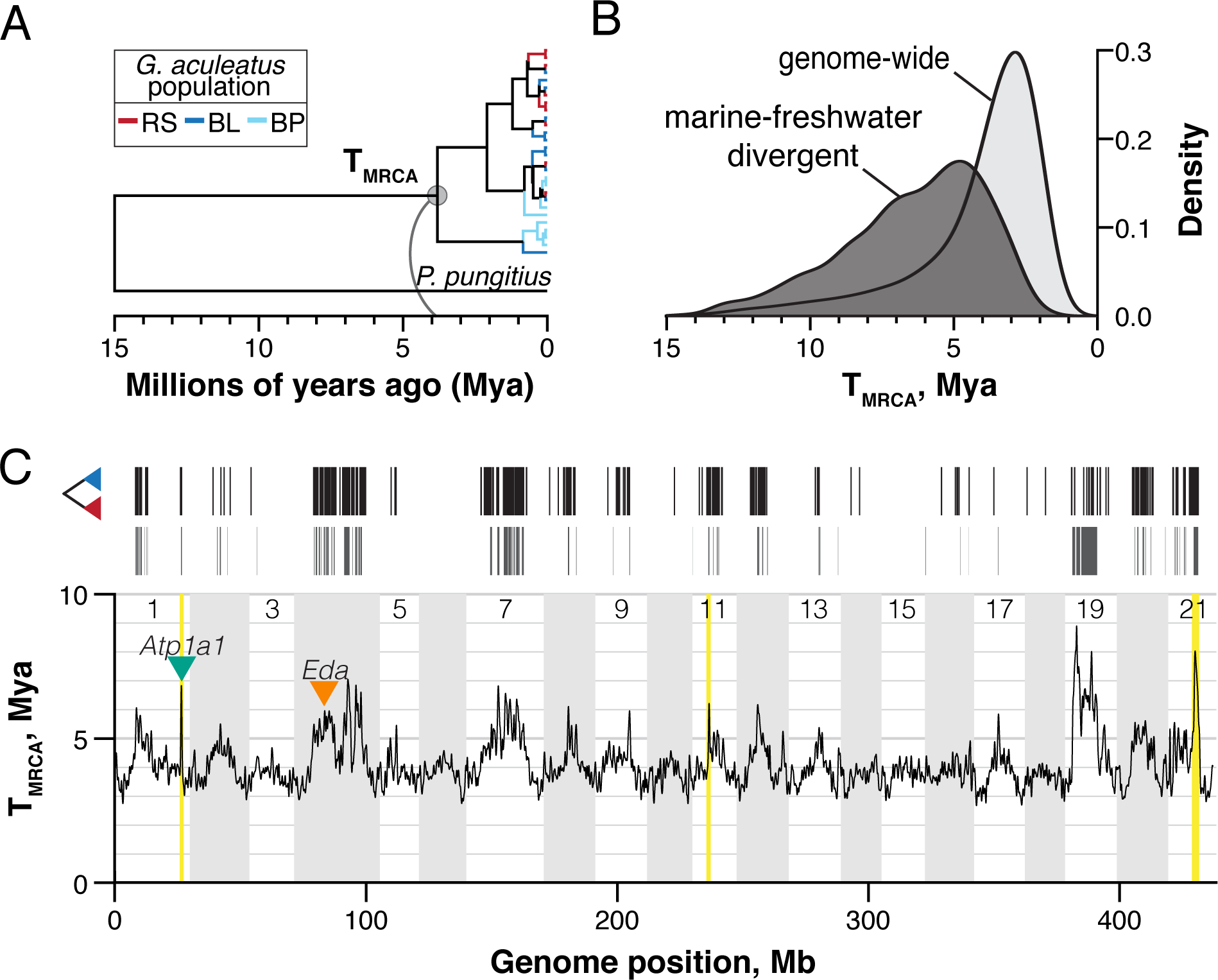
Marine-freshwater divergence has evolved over millions of years, affecting large genomic regions. We performed Bayesian estimation of the time to the most recent common ancestor (T_MRCA_) of alleles at threespine stickleback RAD loci. We calibrated coalescence times within threespine stickleback by including a *de novo* genome assembly from the ninespine stickleback (*Pungitius pungitius*) and setting threespine-ninespine divergence at 15 million years ago. A) Maximum clade credibility RAD gene tree representative of the genome-wide average T_MRCA_. Branches within threespine are colored by population of origin. B) Kernel-smoothed densities of T_MRCA_ distributions for all RAD loci containing a monophyletic group of threespine stickleback alleles (light gray) and those structured into reciprocally monophyletic marine and freshwater haplogroups. C) The genomic distribution of reciprocally monophyletic RAD loci (black, as in Figure 2) is associated with increased T_MRCA_ at a genomic scale. T_MRCA_ outlier windows (those exceeding 99.9% of permuted genomic windows) are shown as gray bars. Genome-wide T_MRCA_ was kernel-smoothed using a normally distributed kernel with a window size of 500 kb. Inverted triangles indicate the locations of *Eda* and *Atp1a1*. Three chromosomal inversions are highlighted in yellow.

We find that the divergence of key marine and freshwater haplotypes has been ongoing for millions of years and extends back to the split with the ninespine stickleback lineage (Fig. 4B). Genome-wide variation averaged 4.1 MY old, and T_MRCA_ for the vast majority of RAD loci was under 5 MY old. In contrast, divergence times at habitat-associated loci averaged 6.4 MYA and, amazingly, the most ancient 10% (118 of 1129) are each estimated at over 10 MY old. This deep genomic divergence not only underscores that local adaptation to marine and freshwater habitats has been occurring throughout the history of the threespine stickleback lineage, for which there is evidence in the fossil record going back 10 million years (Bell *et al.* 1985), but it also demonstrates that at least some of the variation fueling those ancient events has persisted until the present day. In some genomic regions, then, marine and freshwater threespine stickleback are as divergent as threespine and ninespine stickleback, which are classified into separate genera.

Adaptive divergence has impacted the history of the stickleback genome as a whole (Fig. 4C). We identified 32.6 Mb, or 7.5% of the genome, as having elevated T_MRCA_ (gray boxes in Fig. 4C; two-sided permutation test of smoothed genomic intervals, p ≤ 0.001). Outside of the non-recombining portion of the sex chromosome (chr. 19), the oldest regions of the stickleback genome were those enriched for divergent loci. Patterns of ancient ancestry closely mirrored recent divergence in allele frequencies (Fig. 2A) and it appears that historical and contemporary marine-freshwater divergence has impacted ancestry across much of the length of some chromosomes. Chromosome 4, for example, contains at least three broad peaks in T_MRCA_ and a total of 5.9 Mb identified as genome-wide outliers (two-sided permutation test, p ≤ 0.001). This chromosome has been of particular interest because of its association with a number of phenotypes (Colosimo *et al.* 2004; Miller *et al.* 2014), including fitness (Barrett *et al.* 2008). We found the major-effect armor plate locus *eda* comprised a local peak (mean T_MRCA_ = 6.4 MYA) nested within a large region of deep ancestry spanning 8.1 Mb. Moreover, at least two other peaks distal to *eda,* centered at 21.4 Mb and 26.6 Mb, were also several million years older than the genomic average at 6.8 MYA and 7.0 MYA, respectively.

### Long-term divergence maintains linked variation and promotes genomic structural evolution

Intriguingly, genomic regions of elevated T_MRCA_ remained outliers even after removing marine-freshwater relative divergence outlier loci (as measured by F_ST_: Suppl. Fig. S5). We estimated that 7.5% of the genome had increased T_MRCA_ even though only 1.9% of RAD loci (1129 of 57,992) were classified as divergent based on marine-freshwater reciprocal monophyly. When we removed these loci, along with loci with elevated marine-freshwater F_ST_ (F_ST_ > 0.5), many of the regions in which they resided were still T_MRCA_ outliers. It is possible that the remainder of this old variation is neutral with respect to fitness. However, we identified divergence outliers based on only a single axis of divergence: the marine-freshwater axis. Throughout the entire species range, populations are locally experiencing multiple axes of divergence, including lake-stream and benthic-limnetic axes (McKinnon & Rundle 2002), that often shares a common genomic architecture (Deagle *et al.* 2012; Roesti *et al.* 2015). Our data may indicate underlying similarities in selection regimes. Alternatively, this co-localized ancient variation may represent the accumulation of adaptive divergence along multiple axes in the same genomic regions, whether or not the underlying adaptive variants are the same. Aspects of the genomic architecture, such as gene density or local recombination rates, may in part govern where in the genome adaptive divergence can occur (Roesti *et al.* 2013; Aeschbacher *et al.* 2017; Samuk *et al.* 2017). Multiple axes of divergence may therefore act synergistically to maintain genomic variation across the stickleback metapopulation.

Nevertheless, much of the ancient variation we observe may in fact itself be neutral, having been maintained by close linkage to loci under divergent selection between the marine and freshwater habitats (Charlesworth *et al.* 1997). Indeed, the broadest peaks of T_MRCA_ we observe occur in genomic regions with low rates of recombination (Roesti *et al.* 2013; Glazer *et al.* 2015) in other stickleback populations, which would extend the size of the linked region affected by divergent selection. On ecological timescales, low recombination rates in stickleback are thought to promote divergence by making locally adapted genomic regions resistant to gene flow (Roesti *et al.* 2013). Our results potentially extend the inferred impact of recombination rate variation on genomic variation to timescales that are 1000-fold longer, maintaining both multimillion-year-old adaptive variation and large stores of linked genetic variation. Future modeling efforts will be needed to explore the range of population genetic parameter values (e.g. selection coefficients, migration rates, and recombination rates) required to produce the extent of divergence we see here.

Lastly, our findings demonstrate that known chromosomal inversions maintain globally distributed, multilocus haplotypes. The three chromosomal inversions known to be associated with marine-freshwater divergence (Jones *et al.* 2012; Roesti *et al.* 2015) (*inv1*, *inv11*, and *inv21*; yellow bars in Fig. 4C) all showed sharp spikes in T_MRCA_. Genomic signatures of these inversions are distributed throughout the species range, including coastal marine-freshwater population pairs in the Pacific and Atlantic basins (Jones *et al.* 2012) and inland lake-stream pairs in Switzerland (Roesti *et al.* 2015). Despite our limited geographic sampling, our finding that all three of these inversions are over six million years old is further evidence of single, ancient origins of each, followed by their spread across the species range. Each inversion contained a high density of divergent RAD loci (*inv1*: 64% of loci divergent; *inv11*: 60%; *inv21*: 71%) but we also identified regions within these inversions in which haplotypes from marine or freshwater habitats, or both, were not monophyletic. *inv1* and *inv11* both contained two regions separated by loci in which neither habitat type was monophyletic; *inv21*, the largest of the three, contained ten such regions. Additionally, T_MRCA_ and F_ST_ decreased sharply to background levels outside of the inversions, demonstrating the potential for gene flow and recombination to homogenize variation in these regions. We interpret this as evidence that these inversions help maintain linkage disequilibrium among multiple divergently adaptive variants in regions susceptible to homogenization (Kirkpatrick & Barton 2006; Guerrero *et al.* 2012). The presence of these inversions in addition to divergence in regions of generally low recombination (Glazer *et al.* 2015), therefore, further supports the hypothesis that the recombinational landscape can influence where in the genome adaptive divergence can occur (Roesti *et al.* 2013; Samuk *et al.* 2017) and emphasizes the degree to which gene flow among divergently adapted stickleback populations has impacted global genomic diversity.

### Conclusions

Selection operating on two very different timescales — the ecological and the geological — has shaped genomic patterns of SGV in the threespine stickleback. On ecological timescales, selection drives phenotypic divergence in decades or millennia by sorting SGV across geography and throughout the genome (Hendry *et al.* 2002; Hohenlohe *et al.* 2010; Lescak *et al.* 2015; Roesti *et al.* 2015). Our findings show that persistent ecological diversity and continual local adaptation of stickleback has set the stage for long-term divergent selection and for the accumulation and maintenance of adaptive variation over geological timescales. Some of the genetic variants fueling contemporary, rapid adaptation may even have been present – and under selection – since before the threespine-ninespine stickleback lineages split. The genomic architecture of ecological adaptation in one focal population is therefore the product of millions of years of evolution taking place in multiple populations, many of which are now extinct. These findings underscore the need to understand macroevolutionary patterns when studying microevolutionary processes, and vice versa.

## ACKNOWLEDGEMENTS

We thank P. Phillips, M. Streisfeld, J. Postlethwait, K. Sterner for valuable input and lively discussion throughout this project. We also thank K. Alligood, E. Beck, S. Bassham, M. Chase, M. Currey, M. Hahn, L. Fishman, P. Ralph, C. Small, S. Stankowski, J. Willis, two anonymous reviewers, and members of the Cresko Lab and the Institute of Ecology and Evolution for advice and comments on previous versions of this manuscript. J. Postlethwait graciously donated ninespine stickleback tissue, collected under award XXXXXXXX. We acknowledge National Science Foundation awards NSF DEB 1501423 (WAC and TCN), NSF DEB 0949053 (WAC), and National Institutes of Health award NIH T32GM007413 (TCN).

## AUTHOR CONTRIBUTIONS

TCN and WAC conceived of the project and designed sampling, sequencing, and analysis. TCN prepared sequencing libraries, wrote software, and performed data analysis. TCN and WAC wrote the paper.

---

**Figure S1.**
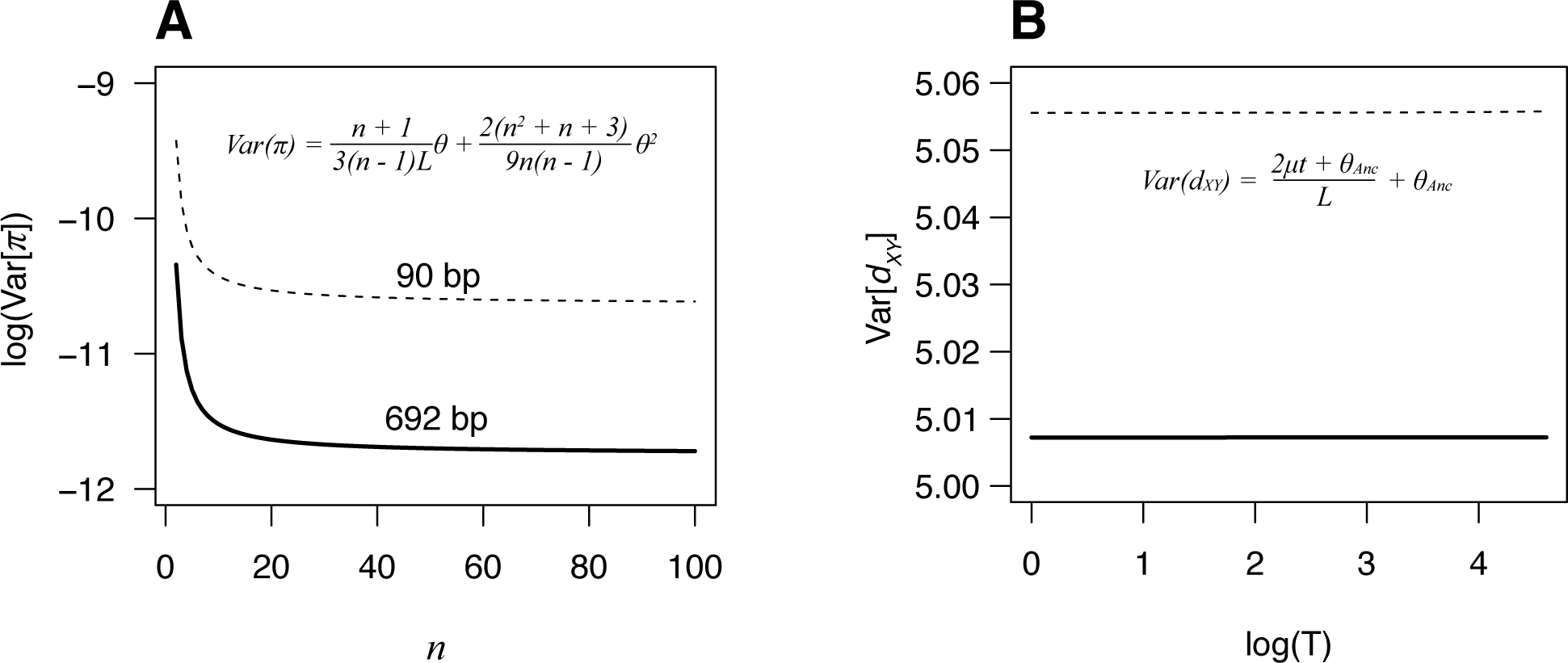
Longer sequences reduce variance in estimates of sequence diversity and divergence. A: Variance in π as a function of the number of chromosomes sampled, using sequence lengths typical of RAD-seq experiments (90 bp) and those in this study (692 bp). Variance was calculated using equation 10.9 in Nei (1987). Right: Variance in d_XY_ (using the equation in box 1 in Cruickshank and Hahn (2014)) as a function of (log-scaled) divergence time of two populations. The change in variance as function of divergence time is dwarfed by the difference in variance obtained with different sequence lengths. In both panels, *n* = sequences sampled; *L* = length of sequence sampled; *θ* = *4Nµ*; *θ*_*Anc*_ = *θ* in the population ancestral to those sampled; *µ* = mutation rate per nucleotide; *t* = time since population split.

**Figure S2.**
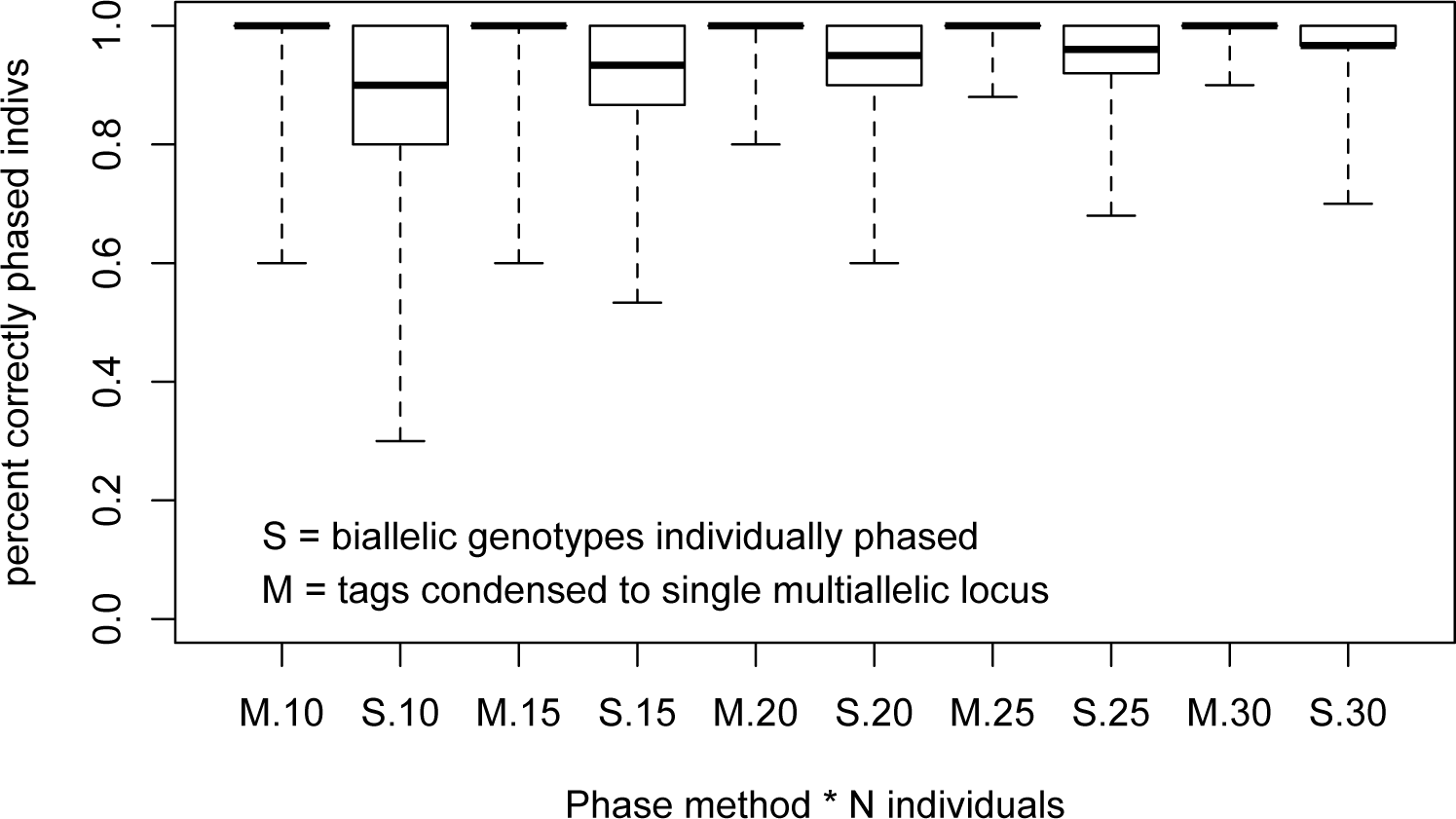
Accurate phasing of RAD loci even at low population-level sampling. Neutrally evolving, non-recombining RAD loci were simulated with ms and seq-gen to generate alignments with of 20 to 60 haplotypes (10-30 diploid individuals) and four to 30 segregating sites. Simulated haplotypes were then ‘cut’ at their midpoints and phased either by inputting all variable sites individually (biallelic ‘SNPs’, S) or by inputting the haplotype information on either site of the cut as multiallelic loci (M). Boxes represent interquartile range (IQR). Bold lines are medians. Whiskers extend to minimum and maximum values. Even with smaller sample sizes (10-15 individuals), over 75% of phasing attempts resulted in 100% phasing accuracy.

**Supplementary Figure S3.**
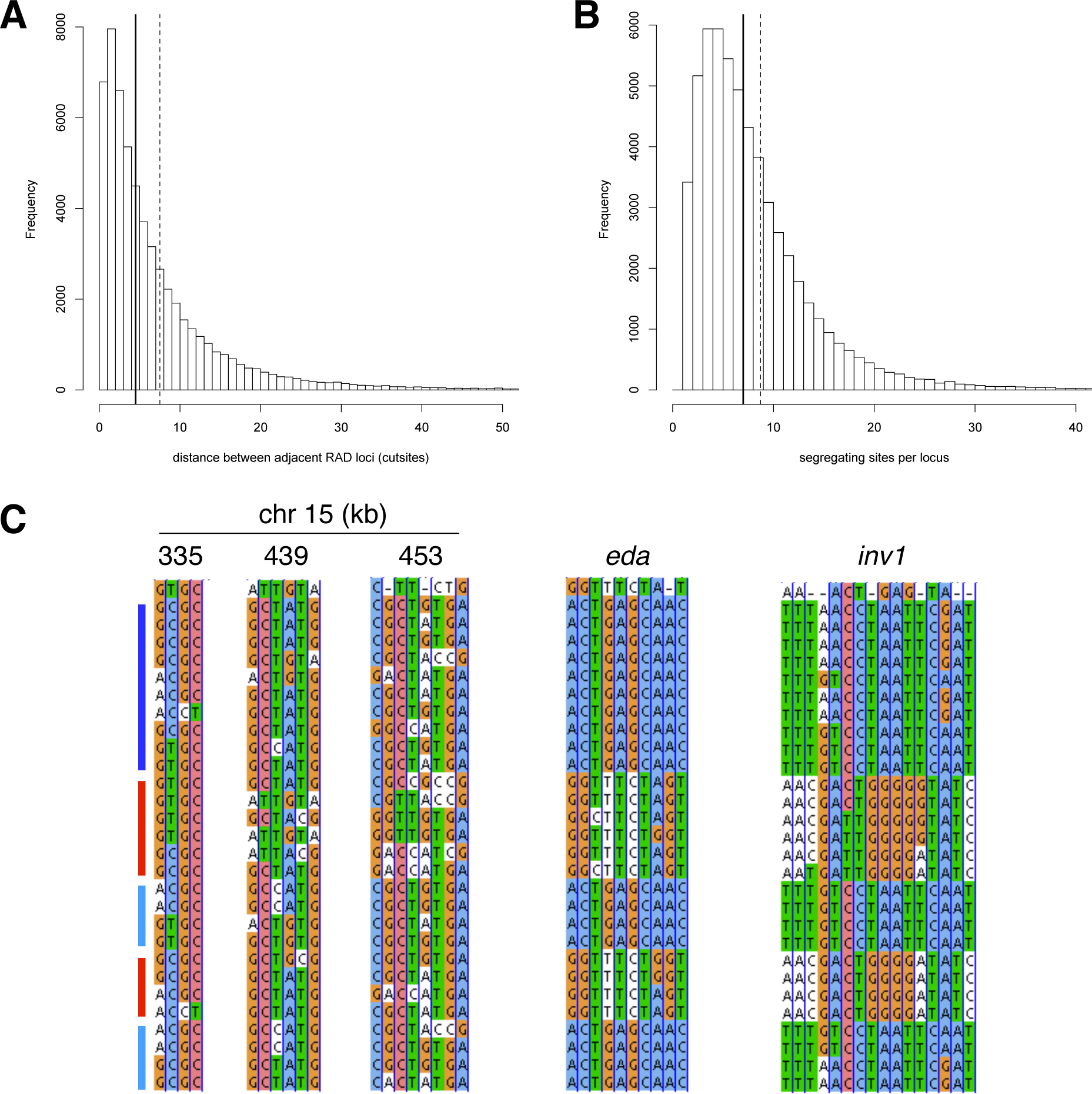
RAD-seq effectively samples genome-wide sequence diversity. Histograms of (A) the distance between adjacent RAD loci (calculated as the distance between the centers of each restriction site) and (B) the number of variable sites per locus show that most RAD loci were within 4 kb of their nearest neighbor and contained ≥ 7 variable sites. Means for each metric are shown as dashed vertical lines. Medians are solid lines. Each histogram is truncated to highlight the bulk of the distribution. Maximum values: distance = 455 kb; variable sites = 155. C: Example haplotypes from five RAD loci in non-divergent (chromosome 15) and divergent (*eda*, *inv1*) genomic regions. Chromosome 15 loci are labeled by their genomic position. The *eda* RAD locus is within the transcribed region of *eda* and *inv1* is within the breakpoints of the chromosome 1 inversion. Colored bars identify population/ecotype of origin. Red: RS (marine); dark blue: BL (freshwater); light blue: BP (freshwater). Alignments visualized in JalView v1.0 (Waterhouse, *et al*, 2009). Only sites that are variable within threespine stickleback are shown.

**Supplementary Figure S4.**
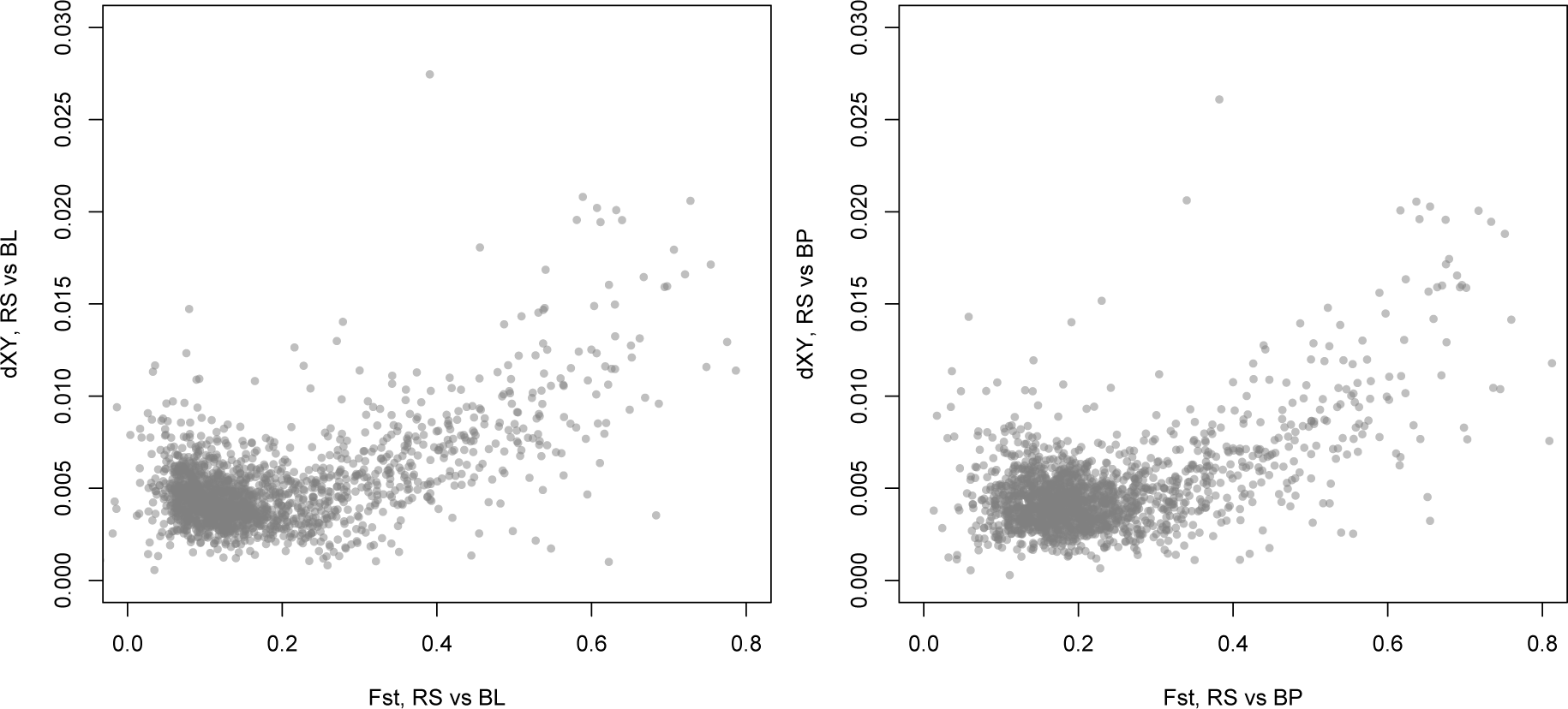
Relative (F_ST_) and absolute (*d*_*XY*_) sequence divergence are positively correlated genome-wide in two instances of marine-freshwater divergence. Points are 250 kb non-overlapping genomic windows. Left panel compares the marine Rabbit Slough population (RS) to the freshwater Boot Lake population (BL) (type-II linear model: r^2^ = 0.314, permuted p-value [reduced major axis] = 0.01). Right panel compares RS to the freshwater Bear Paw Lake population (BL) (type-II linear model: r^2^ = 0.311, permuted p-value [reduced major axis] = 0.01).

**Supplementary Figure S5.**
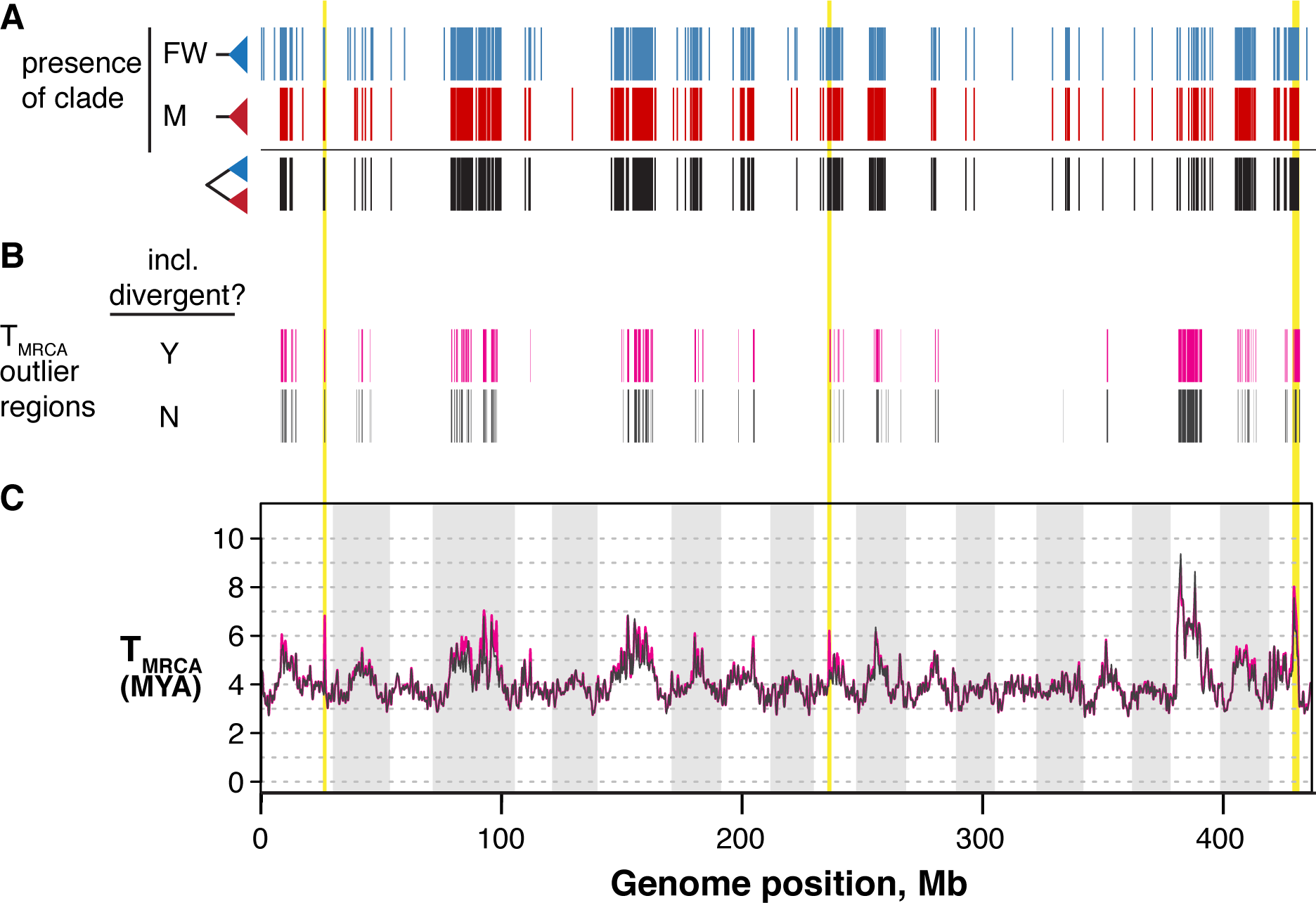
T_MRCA_ outlier regions remain outliers after removing highly differentiated RAD loci. Panel A is taken from Fig. 2 and shows the genomic distribution of reciprocally monophyletic (“divergent”; black bars) RAD loci. Panel B shows the distributions of T_MRCA_ outlier regions (increased T_MRCA_) including all RAD loci (magenta boxes, “Y”). Below are the T_MRCA_ outlier regions after removing divergent loci and any RAD locus with a marine-freshwater (RS vs. [BL+BP]) F_ST_ > 0.5, which is approximately the top 7% of the F_ST_ distribution. Panel C: Genome scans of T_MRCA_ using all RAD loci (magenta) and excluding marine-freshwater outliers (gray).

**Supplementary table 1.**
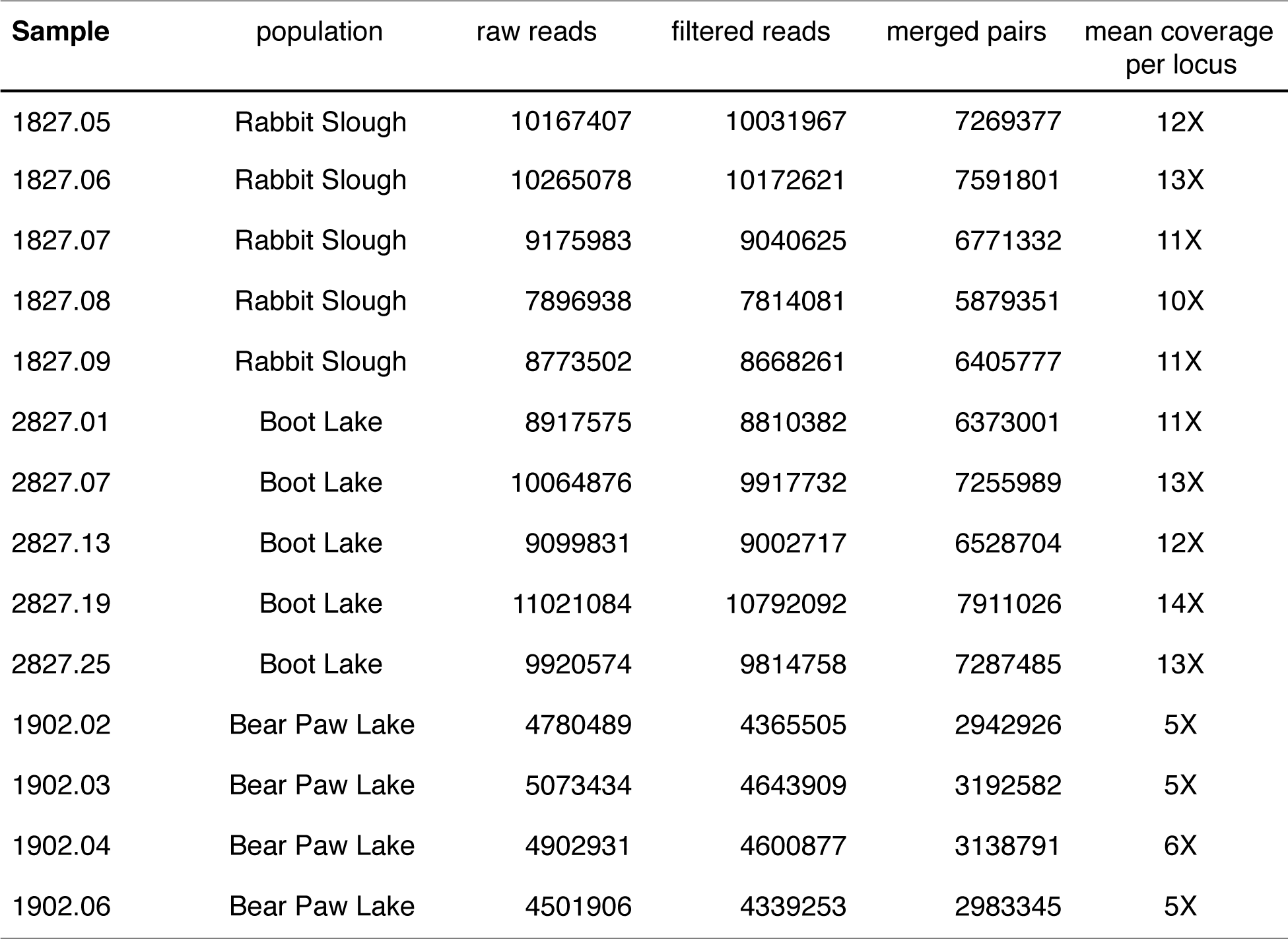
Sequencing summary for threespine stickleback samples

| Sample | population | raw reads | filtered reads | merged pairs | mean coverage<br>per locus |
| --- | --- | --- | --- | --- | --- |
| 1827.05 | Rabbit Slough | 10167407 | 10031967 | 7269377 | 12X |
| 1827.06 | Rabbit Slough | 10265078 | 10172621 | 7591801 | 13X |
| 1827.07 | Rabbit Slough | 9175983 | 9040625 | 6771332 | 11X |
| 1827.08 | Rabbit Slough | 7896938 | 7814081 | 5879351 | 10X |
| 1827.09 | Rabbit Slough | 8773502 | 8668261 | 6405777 | 11X |
| 2827.01 | Boot Lake | 8917575 | 8810382 | 6373001 | 11X |
| 2827.07 | Boot Lake | 10064876 | 9917732 | 7255989 | 13X |
| 2827.13 | Boot Lake | 9099831 | 9002717 | 6528704 | 12X |
| 2827.19 | Boot Lake | 11021084 | 10792092 | 7911026 | 14X |
| 2827.25 | Boot Lake | 9920574 | 9814758 | 7287485 | 13X |
| 1902.02 | Bear Paw Lake | 4780489 | 4365505 | 2942926 | 5X |
| 1902.03 | Bear Paw Lake | 5073434 | 4643909 | 3192582 | 5X |
| 1902.04 | Bear Paw Lake | 4902931 | 4600877 | 3138791 | 6X |
| 1902.06 | Bear Paw Lake | 4501906 | 4339253 | 2983345 | 5X |

**Supplementary table 2.** Genome assembly statistics for *Pungitius pungitius*.

|  | contig (scaffold) |
| --- | --- |
| n | 393,037 (391,396) |
| Max length (bp) | 165,088 (182,644) |
| N50 (bp) | 9,202 (9,886) |
| Average length (bp) | 1,314 (1,320) |
| Gaps (%) | 0.03 |
| Total assembly length (bp) | 516,674,741 |

